# Revisiting the dimensionality of biological diversity

**DOI:** 10.1101/508002

**Authors:** Gabriel Nakamura de Souza, Larissa Oliveira Gonçalves, Leandro da Silva Duarte

## Abstract

Biodiversity can be represented by different dimensions. While many diversity metrics try to capture the variation of these dimensions they also lead to a ‘fragmentation’ of the concept of biodiversity itself. Developing a unified measure that integrates all the dimensions of biodiversity is a theoretical solution for this problem, however, it remains operationally impossible. Alternatively, understanding which dimensions better represent the biodiversity of a set of communities can be a reliable way to integrate the different diversity metrics. Therefore, to achieve a holistic understand of biological diversity, we explore the concept of dimensionality. We define dimensionality of diversity as the number of complementary components of biodiversity, represented by diversity metrics, needed to describe biodiversity in an unambiguously and effective way. We provide a solution that joins two components of dimensionality — correlation and the variation — operationalized through two metrics, respectively: Evenness of Eigenvalues (EE) and Importance Values (IV). Through simulation we show that considering EE and IV together can provide information that is neglected when only EE is considered. We demonstrate how to apply this framework by investigating the dimensionality of South American small mammal communities. Our example evidenced that, for some representations of biological diversity, more attention is needed in the choice of diversity metrics necessary to effectively characterize biodiversity. We conclude by highlighting that this integrated framework provides a better understanding of dimensionality than considering only the correlation component.

## Introduction

Biodiversity encompasses all variation present in life, from genetic material to populations, communities and higher levels of biological organization like entire ecosystems (Wilson 1997). In addition to its broadness in scale and complexity, the central position of the concept of biodiversity in ecological studies justifies efforts to develop measures that properly operationalize the concept. These efforts are reflected in the immensurable number of diversity metrics that have appeared as attempts to encompass all the variation in biodiversity. However, although these diversity metrics allow the description of different dimensions, as the number of them increases the concept of biodiversity becomes operationalized in disparate ways that convey no precise information. This lack of consensus in operationalization of the concept of biodiversity led Hulrbert (1971) to propose the idea of the non-concept of species diversity, in which he advocated that the many metrics of biodiversity be summarized in only a few relevant ones that can be used to express adequately and unambiguously the concept of biodiversity.

Long since Hulrbert’s seminal work, there has been a pronounced increase in the number of metrics that quantify characteristics of biological diversity other than the traditional taxonomic-based metrics, revealing that patterns of diversity for some communities can be best described using other components of biological diversity, such as functional and phylogenetic components (Graham and Fine 2008, Cisneros et al. 2014). However, these findings are not consensual (e.g Lamb et al. 2009), since some phylogenetic and functional metrics can be strongly correlated with traditional metrics (Tucker and Cadotte 2013, Tucker et al. 2018), deepening the question of which metrics represent the fundamental components of biological diversity (Hulrbert, 1971). A theoretical approach to searching for fundamental variation in biodiversity is to integrate the many sources of information in a unique framework. This integration can be achieved by investigating the relationships among existing metrics. A previous work that proposed this integration based it on quantifying a characteristic of biodiversity known as dimensionality (Stevens and Tello 2014).

Dimensionality can be defined, at the community scale of biological organization, as the amount of information needed to effectively characterize the variation presented in a given biodiversity representation, by means of diversity metrics. Communities with high dimensionality require more dimensions to be effectively described than communities with low dimensionality (Stevens and Tello 2014). Quantifying the dimensionality of biodiversity currently involves searching for the degree of complementarity in spatial or temporal variation among multiple metrics of diversity, which is obtained mainly through a measure denominated Eveness of Eigenvalues (hereafter EE) (Stevens and Tello 2014).

Stevens and Tello’s EE metric is obtained by Principal Component Analysis (PCA) of a matrix of diversity metrics (hereafter matrix **M**, *sensu* Ricotta 2005) for a set of communities, and calculating an evenness metric for the eigenvalues of the axes that represent this fundamental biodiversity space. The logic behind EE is that, if the diversity metrics used to characterize communities have low complementarity, almost all of the fundamental variation in biodiversity will be concentered in a few axes, producing a low EE. On the other hand, if diversity metrics are completely complementary with each other (variation in biodiversity will be equally distributed among axes) the EE of the communities will be 1.

The EE metric represents, in a simple way, the degree of complementarity among the dimensions of biodiversity represented by diversity metrics, which comprises what we call here the correlation component of dimensionality (see also Tucker and Cadotte 2013, Lamb et al. 2006 for uses of correlation component). However, EE ignores another source of information in dimensionality — the amount of variation, or importance, that each diversity metric presents in fundamental biodiversity space. This comprises what we call here the variation component of dimensionality.

Suppose a situation in which diversity metrics are highly correlated (Figure 1A) and each metric accounts for a similar amount of variation in fundamental biodiversity space (Figure 1B). This situation has low complementarity among dimensions of biodiversity and high redundancy in the amount of variation that each metric captures in fundamental biodiversity space (represented as the length of the arrows in 1B). Consequently, we could rely on any of these diversity metrics to effectively represent the variation in biodiversity of these communities. On the other hand, communities with low complementarity may present a situation in which one of the metrics captures almost all the variation in the fundamental biodiversity space (Metric 2 in Figure 1C), indicating low redundancy of metrics. Following the current approach to measuring dimensionality, EE would indicate similar patterns of dimensionality for communities in 1B and 1C. However, the choice of metric in 1C is of greater importance than in 1B, in which the metrics are highly redundant regarding the information captured. Therefore, considering only the correlation component does not provide enough evidence to support the decision of which diversity metrics to use to effectively characterize biological diversity for two communities with similar EE, because it disregards the variation component inherent to dimensionality.

**Figure 1:**
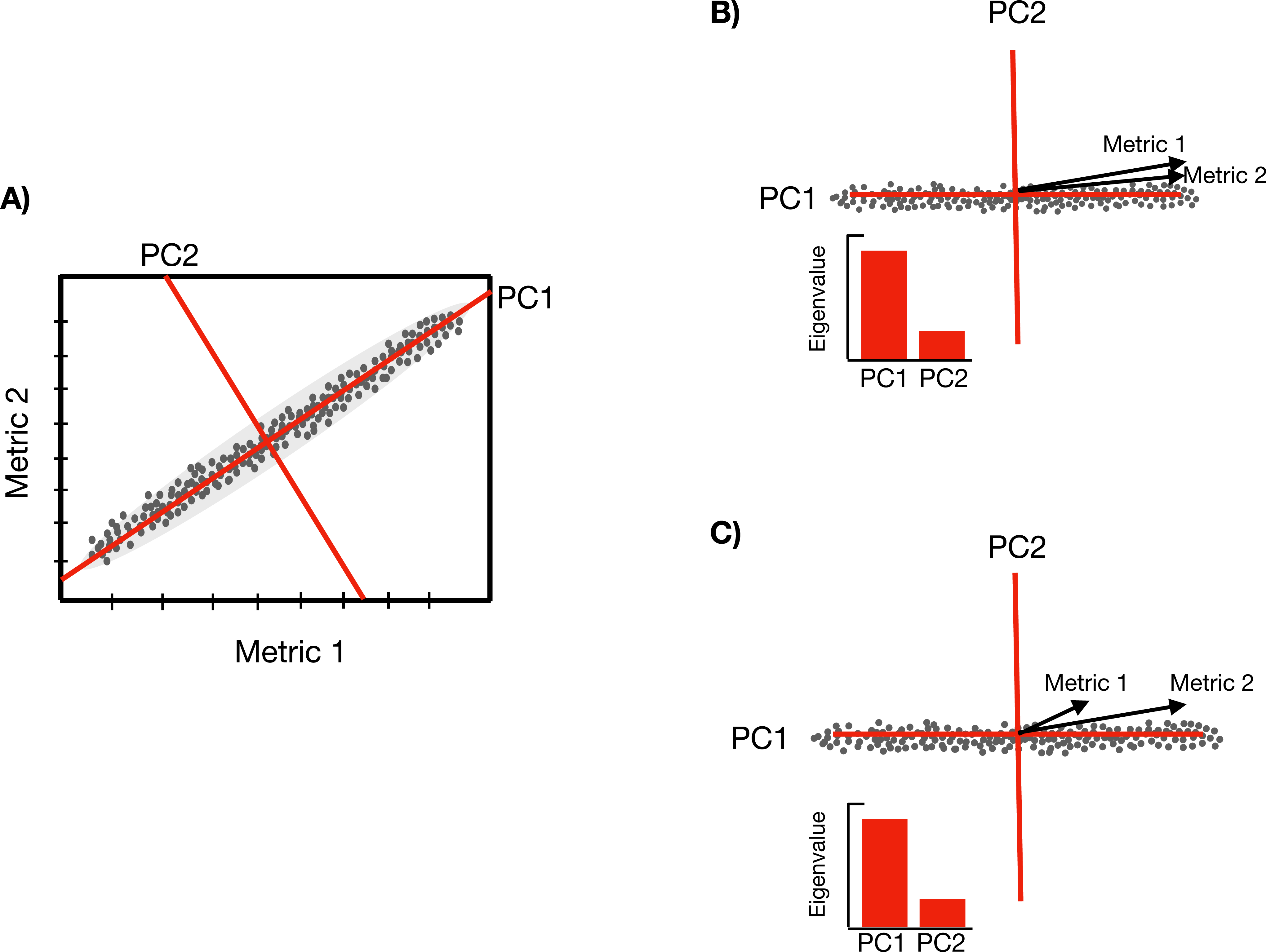
A) A set of communities described by two diversity metrics (Metric 1 and Metric 2) that are highly correlated. This pattern of correlation can be related to two diversity metrics that account for similar amounts of variation in the reduced biodiversity space (B, high redundancy), or be a situation in which one metric has disproportional importance for capturing variation in biodiversity space (C, low redundancy).

Finding a measure that captures the variation component of dimensionality is not an impediment for effectively characterizing dimensionality, since it can be operationalized by the metric Importance Values (hereafter IV) proposed by Wilsey et al. (2005). However, since the common way to quantify dimensionality (Stevens and Tello 2014) is limited to capturing only the correlation component, the development of a unified framework that combines both correlation and variation components would provide a way to better represent the dimensionality of biodiversity.

Therefore, our aim was to update the concept of dimensionality of biodiversity and its operationalization by integrating the correlation and variation components through EE and IV in a framework for quantification of dimensionality. To do this we show, through simulation, how EE and IV together can distinguish situations with different degrees of complementarity of dimensions of diversity and redundancy of information that each metric captures. We then present an empirical example of the investigation of dimensionality by applying the integrated framework to communities of small mammals (cricetids and marsupials). Specifically, we evaluated the level of complementarity and redundancy for different sets of diversity metrics used to describe the biodiversity of cricetids and marsupials, highlighting how the proposed dimensionality framework facilitates the first step of biological assessment — the choice of metrics to be used for characterizing biodiversity.

## Material and Methods

### Investigating the dimensionality of biodiversity: obtaining EE and IV

Our framework for investigating the dimensionality of biodiversity comprises three steps. The first step is to calculate matrix **M**, which, for the sake of simplicity, will contain three metrics of diversity for the simulation analysis: a measure of functional diversity (FD [Petchey & Gaston 2006)]), a measure of phylogenetic diversity (PD [Faith 1992]) and richness. We chose a simplistic approach with only three metrics since our objective with the simulation analysis was to focus on showing how IV can reveal patterns that are not detected by using only EE. We were more interested in the patterns of correlation and variation of diversity metrics in biodiversity space than the particularity of the metrics themselves. We present a more realistic exploration of the integrated framework in the section *Assessing the dimensionality of biodiversity in small mammal communities.*

The second step involves performing a PCA of matrix **M** using a standardized correlation matrix. As will be shown next, the standardization method applied to matrix **M** prior to the PCA must differ between the calculation of EE and IV.

The third step is to calculate the dimensionality metrics EE and IV. We calculate EE using Camargo’s evenness index in Equation 1, following the original proposition of Stevens and Tello (2014):

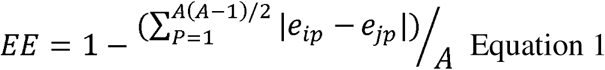

Camargo’s evenness index (Camargo 1993) is calculated using the axes (A) and their respective eigenvalues (*e*_*ih*_ and *e*_*jh*_) from a PCA of the standardized matrix **M**, in which the metrics were scaled to have a mean of zero and equal variances. The higher the value of EE, the higher the complementarity the communities have in relation to the dimensions of biodiversity represented in matrix **M**. On the other hand, lower EE values indicate lower complementarity in the dimensions used to characterize the communities. IV is calculated according to the method proposed by Wilsey et al. (2005), using a matrix (**M**) standardized by the maximum values of each diversity metric. This standardization removes the effect that the different units of each diversity metric have, without modifying their original variation. To obtain IV for each diversity metric in matrix **M** we apply Equation 2, in which IV_i_ represents the IV of diversity metric *i*, r^2^_ij_ is the squared correlation of diversity metric *i* with PC_*j*,_ and R^2^_j_ is the amount of variation that PC_j_ accounts for in ordination space (biodiversity space).

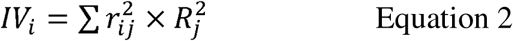

PC varies from 1 to *j* and corresponds to the number of significant eigenvectors in the PCA, evaluated by the Kaiser-Gutmann criterion. The greater the IV the more variation the diversity metric accounts for in biodiversity space. IV approaches 1 when the diversity metric accounts for almost all the variation and approaches zero when the metric accounts for little variation. Sets of communities with highly uneven IV values for diversity metrics possess low redundancy in metric importance, while communities with highly even IV values possess high redundancy regarding the amount of information captured by each metric.

### Testing the assessment of the dimensionality of diversity using EE and IV

To assess the effectiveness of EE and IV in acquiring information regarding correlation and variation of dimensionality in matrix **M**, the following conditions must be met: (1) EE values must not differ for set of communites simulated in scenarios with the same level of correlation among diversity metrics, and must differ among communities that have different levels of correlation among diversity metrics; (2) for scenarios with low and high correlation, IV must be similar among metrics that have similar variation in biodiversity space (e.g. Figure 1B), and differ for scenarios in which variation in biodiversity space is mainly due to a single metric (e.g. situation represented Figure 1C, Metric 2 must have a higher IV than Metric 1). We evaluate whether EE and IV can recover these patterns by simulating communities with varying degrees of correlation and variation for each metric in biodiversity space obtained from matrix **M**.

The simulations were based on a pattern-oriented procedure, producing diversity metrics with patterns of correlation and variation that represent four scenarios with the following characteristics: In the HiC/EqV (High Correlation and Equal Variation) scenario the diversity metrics are highly correlated and have similar variation in biodiversity space. The HiC/DifV (High Correlation and Different Variation) scenario has diversity metrics that are highly correlated and vary in importance of each metric in biodiversity space. The LoC/EqV (Low Correlation and Equal Variation) scenario has diversity metrics with low correlation and similar importance in biodiversity space. Finally, the LoC/DifV (Low Correlation and Different Variation) scenario has diversity metrics with low correlation and dissimilar importance in biodiversity space.

We generate scenarios HiC/EqV and HiC/DifV by starting with a phylogeny that was simulated by a birth-death processes (function *sim*.*bdtree* from the package geiger [Harmon, Weir, Brock, Glor, & Challenger, 2008]) where a species, chosen randomly, initiates the procedure by colonizing a given community. Subsequent addition of species to the community depends on the species that are already present in that community. Communities at one extreme will only contain species that are phylogenetically closely related to each other (top 10%), with the phylogenetic filter becoming less restrictive until communities do not have any phylogenetic filter that restricts coexistence of species (least restrictive condition). Since we simulated a continuous trait that was conserved over the phylogenetic tree — evolved according to a Brownian motion model, using the function *rTraitCont* (Paradis et al. 2004) with the ρ [*rho*] parameter set to 3 — with the number of species in each community gradually increasing (less phylogenetic filter, more species), the procedure created a gradient of phylogenetic, functional and taxonomic diversity metrics. In order to generate differences in variation of the diversity metrics, in scenario HiC/ DifV we simulated a trait that evolves according to a regime of stabilizing selection (Ornstein-Uhlebeck model with the strength of selection set by the parameter α at 0.8) that restricts trait variation to within an optimal range (represented by a θ [theta] of 0). This allowed us to generate a set of communities in which the diversity metrics were highly correlated but variation of FD was much lower than that of richness and PD since the traits that were used in the calculation of FD were restricted by the selection process.

We generated the scenario LoC/DifV by following the same procedures described above for scenario HiC/EqV, however, the trait was simulated to have low phylogenetic signal and the phylogenetic tree used to calculate PD was modified to simulate a process of evolution in which most speciation occurs near the root (a star-like phylogeny). This procedure resulted in low correlation between PD and FD, since the relationship between phylogeny and traits was disrupted. Additionally, low variability for PD and richness metrics was obtained since we set the simulations to produce communities with the same number of species but with the phylogenetic filtering acting in community assembly. Consequently, most of the variation in this scenario is due to the FD metric. Finally, to generate scenario LoC/EqV we simulated communities in which all species in the phylogenetic tree had an equal probability of occurring in any community (no phylogenetic filtering acting on the assembly), and set the richness to be very similar for all communities. This procedure generated metacommunities with low correlation and similar amounts of variation for all diversity metrics.

We generated 999 sets of communities for each scenario described above, with the metacommunities of all scenarios being composed of 50 communities with a minimum of 20 and a maximum of 200 species. The phylogenetic filter was set to act gradually on the communities, increasing by the order of 10% (start by selecting the top 10% most phylogenetically similar species, followed by the top 20% and so on until 90% of the species have been selected from the pool). Details and an illustration of the simulation procedures and scenarios are presented in the supplementary material Appendix S1, along with a link to an interactive module that we produced to illustrate the simulation procedure used in this work.

Finally, we tested whether the values of EE and IV met our theoretical expectations. We checked if EE values differed between scenarios with low correlation and scenarios with high correlation (scenarios HiC/DifV and HiC/EqV versus scenarios LoC/EqV and LoC/DifV). To effectively capture the correlation component of dimensionality EE must be higher in scenarios with low correlation among diversity metrics than in scenarios with high correlation. To test for differences among IV values of each metric in the scenarios we used a graphical tool called profile of importance (Wilsey et al. 2005) and quantified differences in IV of each metric by calculating F values obtained from a linear model (Equation 3). F values allow the IV values of the three dimensions (PD, FD and richness) to be compared and to determine if the IV values of the DifV scenarios (scenarios HiC/DifV and LoC/DifV) differed more from each than did the IV values calculated for the EqV scenarios (scenarios HiC/EqV and LoC/EqV). The simulation scenarios and the theoretical expectations regarding EE and IV follow the schematic representation present in Figure 2.

**Figure 2:**
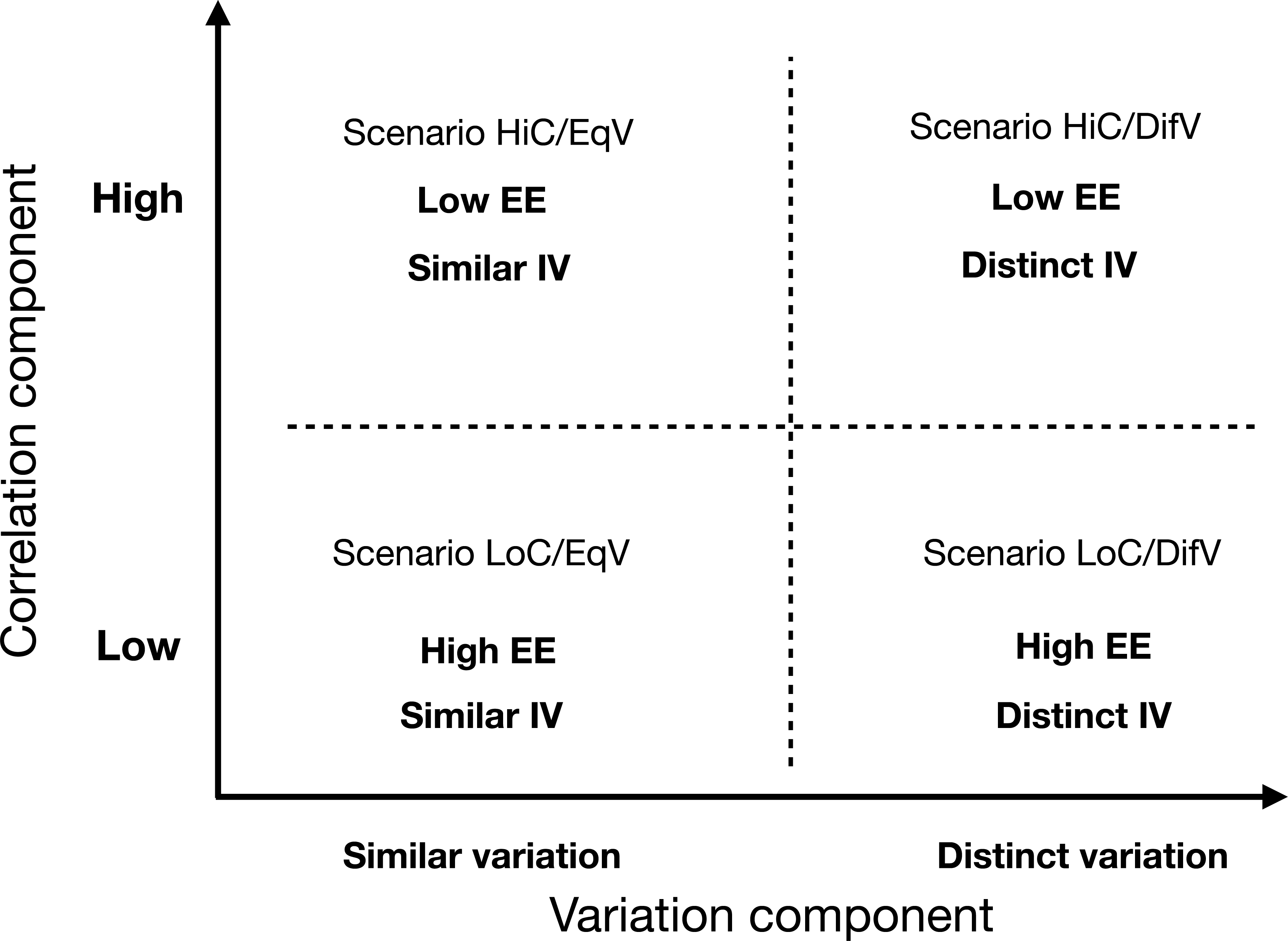
Schematic representation of simulated scenarios and expected outcomes for EE and IV. The abscissa represents the variation component of dimensionality. Metacommunities were simulated to show similar values of variation among metrics (lower left quadrant) or different values of variation among metrics (lower right quadrant), so that, respectively, similar and different IV values among diversity metrics are expected. The ordinate represents the correlation component of dimensionality. Metacommunities were simulated that had metrics with high (upper right panel) and low correlation, so that, respectively, low and high EE values are expected.

### Assessing the dimensionality of biodiversity in small mammal communities

We illustrate the application of the dimensionality framework with a database of small mammal communities (marsupial and cricetid mammals) distributed throughout the South American continent. We constructed matrix **M** for these communities by calculating eight diversity metrics that represent different dimensions of taxonomic, functional and phylogenetic components of biological diversity. The choice of metrics was based on the works of Tucker et al. (2017) and Scheiner (2019), which together represent the most complete compilation and classification of metrics of taxonomic, functional (Scheiner, 2019) and phylogenetic diversity (Tucker et al. 2017). We chose at least one metric for each of the richness, divergence and regularity dimensions of the three components of biodiversity considered here. The taxonomic component was represented by richness; the functional component by FD (richness dimenson, Petchey and Gaston 2006b), FEve (regularity dimension) and FDiv (divergence dimension, Villéger et al. 2008); and the phylogenetic component by PD (richness dimension, Faith 1992), MNTD (divergence dimension, Webb et al., 2002), PSV (divergence dimension, Helmus et al., 2007) and PE_ve_ (regularity dimension, Villéger et al. 2014).

Traits used to calculate functional metrics comprised life-history attributes — weight, head-body length, diet and form of locomotion. Species were categorized according to their diet as insectivores, herbivores, granivores, omnivores, frugivores, piscivores, seed predators and leaf predators, and according to their modes of locomotion as terrestrial, semifossorial, semiaquatic, arboreal and scansorial. Some species were allocated to more than one diet and locomotion category. All calculated diversity metrics require a distance matrix or a functional dendrogram obtained from a distance matrix. Therefore, to obtain the functional distance matrix we used Gower distance (Pavoine et al. 2009) for traits that have different statistical characteristics (numerical and categorical).

The phylogenetic hypothesis used to calculate phylogenetic indices was obtained from the mammalian phylogenies of Bininda-Emonds et al. (2007) and Fabre et al. (2012), the latter of which was used to improve the phylogenetic resolution to species level. Seven species present in our data were not included in the phylogeny Fabre et al. (2012), so we included these species as polytomies within their respective genera. Divergence times for our phylogeny were estimated in millions of years by equally distributing the ages of undated nodes, based on the know ages present in Bininda-Emonds et al. (2007) and Fabre et al. (2012), using the BLADJ algorithm of Phylocom software (Webb et al. 2008). The phylogenetic hypothesis and the original references compiled to assemble the community data used in this work are provided in Figure S2 and Table S1 of Appendix 2 of the supplementary material.

The metrics EE and IV were calculated as previously described, with the number of axes used in IV calculation being determined by the Kaiser-Gutmann stop criterion. We also compared the observed values of EE with a null distribution of 999 EE values generated by a null model that randomizes a species incidence matrix while preserving differences in richness among sites and mixing species frequency (performed with the *sim3* function from the *EcoSimR* package [Gotelli and Ellison 2013a]). Using this null model we tested the null hypothesis that observed EE values do not differ from expected EE values according to variation in richness. We implemented a function called *dimensionality* to calculate EE values from matrix **M**. The function allows the user to choose the evenness method that will be used in the calculation. It can be accessed at https://github.com/GabrielNakamura/dimensionality_function.

We calculated IV for the small-mammal metacommunities according to Equation 2, applying *ImportanceVal* — the R code for the IV function (the function can be accessed at https://github.com/GabrielNakamura/IV_function). We used the Kaiser-Gutmann stop criterion and a bootstrap procedure that re-sampled matrix **M** 999 times and recalculated IV for each metric so that we generated confidence intervals for the IV value of each diversity metric. We performed all calculations with a standardized matrix **M** (scaled to a mean of zero and unit variance for the calculation of EE values and standardized by the maximum values of each metric for the calculation of IV values). Bootstrapped IV values were submitted to an Ordinary Least Square (OLS) linear model to test for differences in the importance of the components of diversity that assemble matrix **M**:

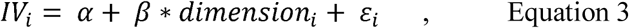

Equation 3 represents the effects parametrization model in which *IV*_*i*_ represents the predicted value of IV for the diversity metric *i,β* the effect of a given dimension over another and *ε*_*i*_ the error term associated with the residuals, which follow a Gaussian distribution. Each value of IV was classified as belonging to the phylogenetic (PD, PEve, PSV and mntd), the functional (FD, FDis and FDiv) or the taxonomic (richness) dimension. Through this model we aimed to determine if any of the components of diversity (functional, phylogenetic or taxonomic) captures a greater amount of information from biodiversity space. Additionally, we performed another linear OLS model using the same set of data but considering each metric as the explanatory variable, in order to assess differences in importance among diversity metrics. For both models we performed a Tukey test to assess pairwise differences in importance among dimensions and metrics.

The dimensionality framework was applied to four different configurations of matrix **M**: all metrics; a combination of phylogenetic metrics and richness; a combination of functional metrics and richness; and a combination of functional and phylogenetic metrics. We performed these analyses to show how dimensionality can change depending on the components of diversity used in matrix **M**, and what the implications of different values of EE and different similarities among metrics IV (represented as Camargo’s evenness of IV metrics) are on the choice of diversity metrics to be used to represent the biodiversity. For these analysis we also computed EE as the mean value calculated from a bootstrap procedure equivalent to that used for the IV metric, in order to generate confidence intervals.

## Results

### Simulated data

Our simulation revealed that EE and IV, when used together, acquire information regarding two aspects of dimensionality: correlation among metrics and the variation that each metric accounts for in biodiversity space. This complementary information that IV brings to the analysis of dimensionality is evidenced in Figure 3. Thus, different patterns of redundancy in information captured by the metrics can be obtained for a given level of correlation, with greater differences among IV values in scenarios HiC/DifV and LoC/DifV (right side of Figure 3) than in HiC/EqV and LoC/EqV (left side of Figure 3).

**Figure 3:**
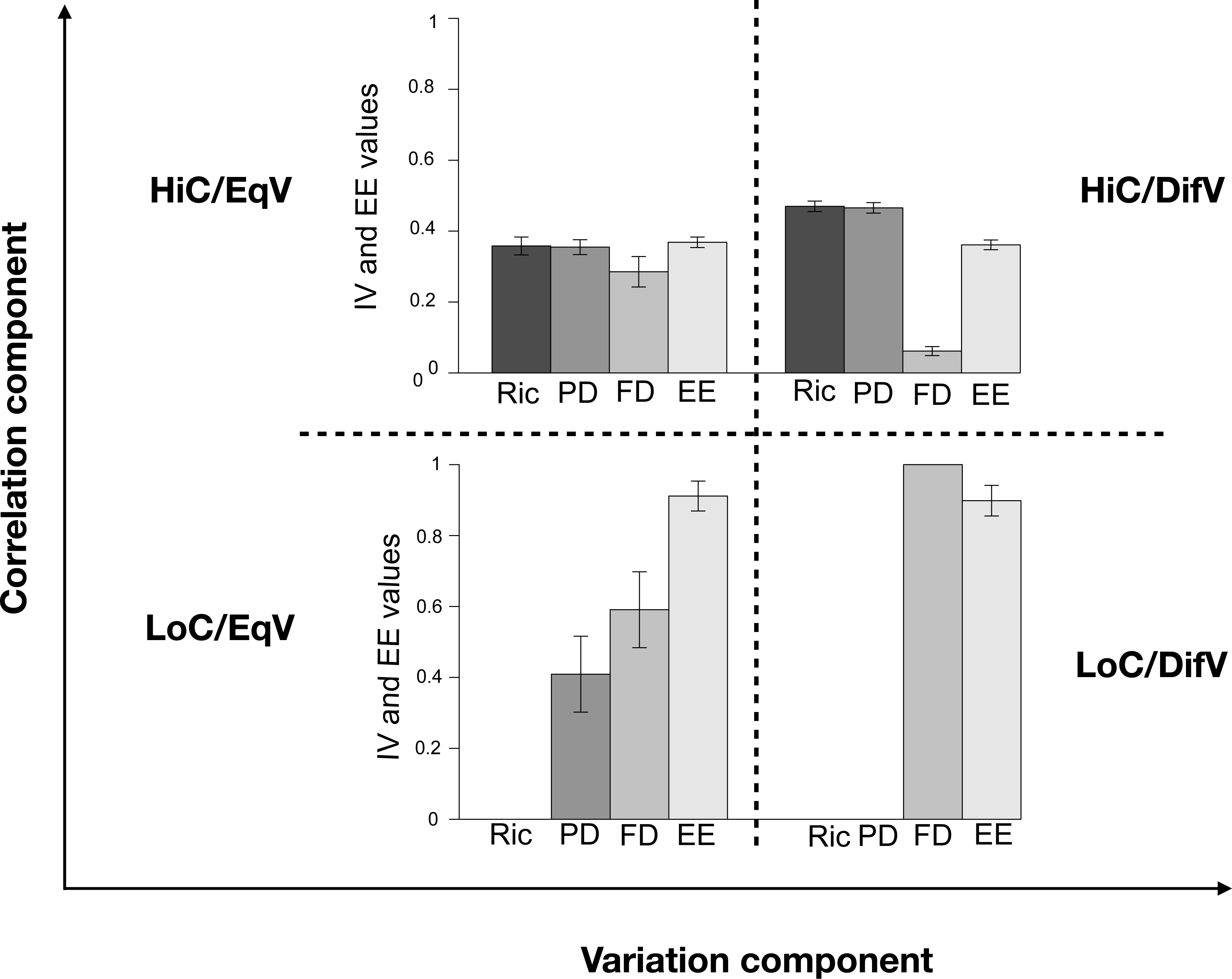
Bar plots showing IV and EE calculated for metacommunities simulated according different scenarios (HiC/EqV, HiC/DifV, LoC/EqV and LoC/DifV) using PD, FD and richness metrics in matrix **M**. For each of these scenarios situations were presented in which the metrics contribute similarly or unequally in biodiversity space (variation in ordinate axis) and are highly or lowly correlated (variation in abscissa axis).

The differences in EE between scenarios of high and low correlation (Figure 1, comparison between EE of upper and lower graphics), but not between scenarios of different and equal variation (Figure 1, comparison between EE bars in the same row) support our argument that this metric captures only the correlation component of dimensionality.

The ability of IV to capture the degree of redundancy in biodiversity information of the metrics was clear mainly for the HiC/DifV scenario, in which the attribute used to generate communities exhibited low variation (OU model) and, consequently, the FD metric presented lower IV than richness and PD metrics. It is worth noting that differences among the IV of metrics was greater in scenario LoC/EqV than in scenario HiC/EqV (Figure 1, lower right graphic), since it is not possible to obtain high redundancy in metric information (indicated by similar IV values among metrics) along with high values of complementarity (indicated by high EE). High redundancy in the importance of metrics is only possible for communities with low EE (low complementarity of dimensions), as demonstrated by scenario HiC/EqV. The magnitude of the differences in IV among metrics for each scenario is shown in Figure S3 of Appendix S3 of the supplementary material.

### Small mammal communities

We obtained a moderate value for complementarity for the small mammal communities, as indicated by an EE of 0.49 for matrix **M** calculated with all eight diversity metrics. The correlation component of dimensionality, at least for the three analyzed components of diversity (functional, phylogenetic and taxonomic), may be a consequence of spatial gradients of species richness, as evidenced by comparing observed EE with that expected by the null model distribution of EE (Figure S4 in Appendix 3 of the supplementary material).

Only two axes of the PCA were significant according Kaiser-Guttman criterion (representing 70% of all the variation in matrix **M**), and composed the fundamental biodiversity space in which IV was calculated. Observed IV values for the eight diversity metrics ranged from 0.19 for PSV (27% of all the variation in biodiversity space) to 0.003 to FDiv (0.3% of all the variation in biodiversity space). Bootstrap means and confidence intervals for IV for all metrics are illustrated in Figure 4 through the IV profile (*sensu* Willig and Hollander 1995), evidencing PSV as the metric capturing most of the variation in biodiversity space, followed by richness.

**Figure 4:**
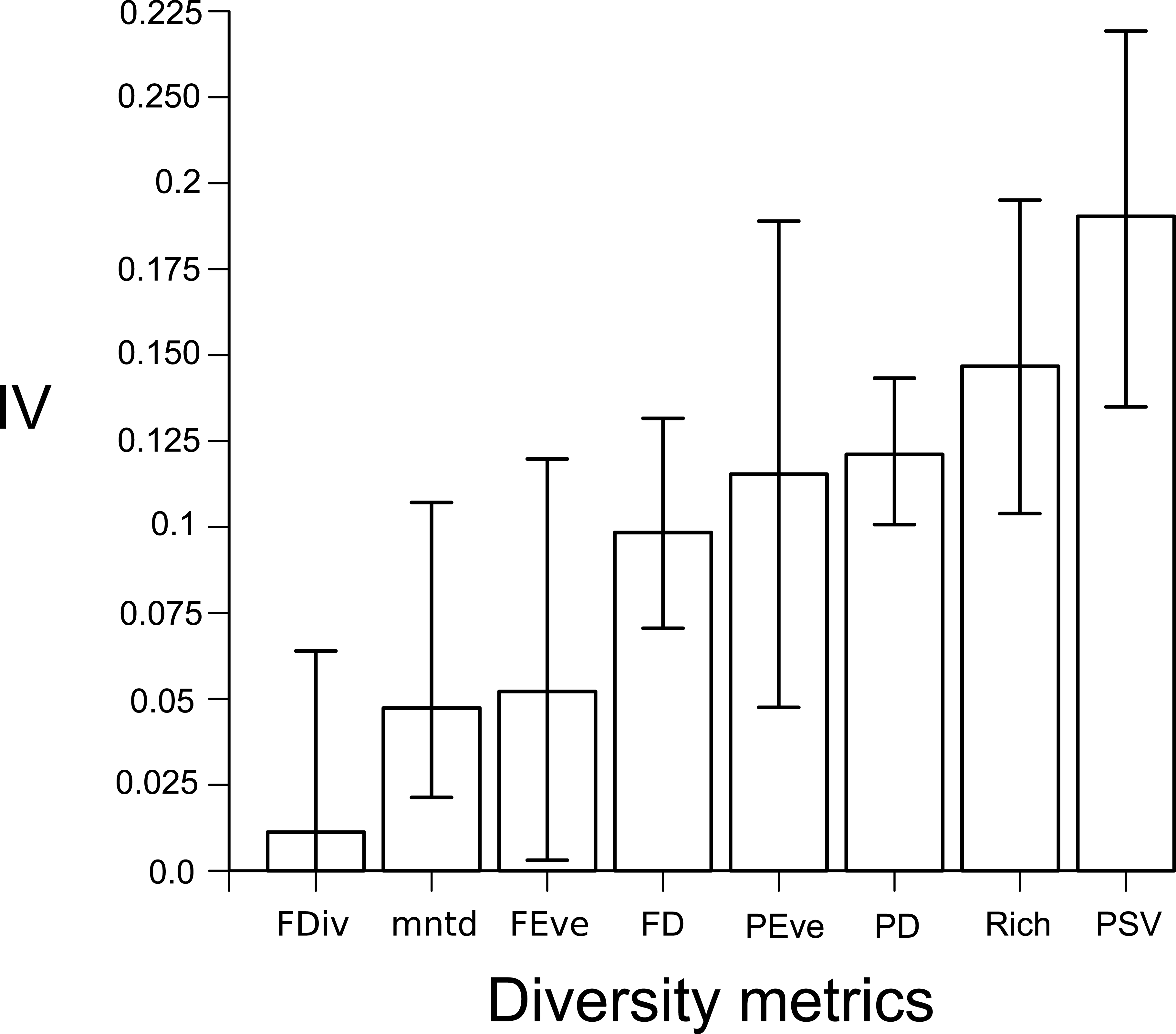
IV profile for marsupial and cricetid mammal communities from South America calculated using matrix **M** containing eight diversity metrics. Bar height corresponds to the mean IV for each diversity metric while lines represent 95% confidence intervals, both calculated via a bootstrap procedure.

The linear OLS model showed significant variation in IV among diversity metrics (F-value= 3.428; p<0.05), while the Tukey test revealed that the greatest difference in importance was between taxonomic and functional components of biodiversity followed by the difference between phylogenetic and functional components (difference between observed means of 0.092 and 0.064, respectively; Figure S5 of Appendix 3). This finding highlights the importance of considering the taxonomic and phylogenetic dimensions in characterizing the biodiversity of communities of cricetids and marsupials.

Analysis of dimensionality for matrix **M** containing functional metrics and richness had the highest complementarity (highest EE) and lowest redundancy in metric importance (biodiversity representation with similar values of IV, as indicated by a lower evenness of IV than obtained for other sets of metrics) (Figure 5). PSV was the metric that captured the most information in matrix **M** containing phylogenetic metrics and richness (30% of all the variation in biodiversity space) and phylogenetic and functional metrics (31% of all the variation in biodiversity space), as well as for matrix **M** containing all metrics (24% and of all the variation in biodiversity space). For matrix **M** that considered only functional metrics and richness, richness captured most of variation (47% of all the variation in biodiversity space). Despite the high variability, as indicated by the confidence intervals of IV and EE evenness, it is worth noting that IV evenness remains constant for different mean values of EE, with the greatest IV evenness being for the set of metrics that had the lowest EE value (matrix **M** with phylogenetic metrics and richness).

**Figure 5:**
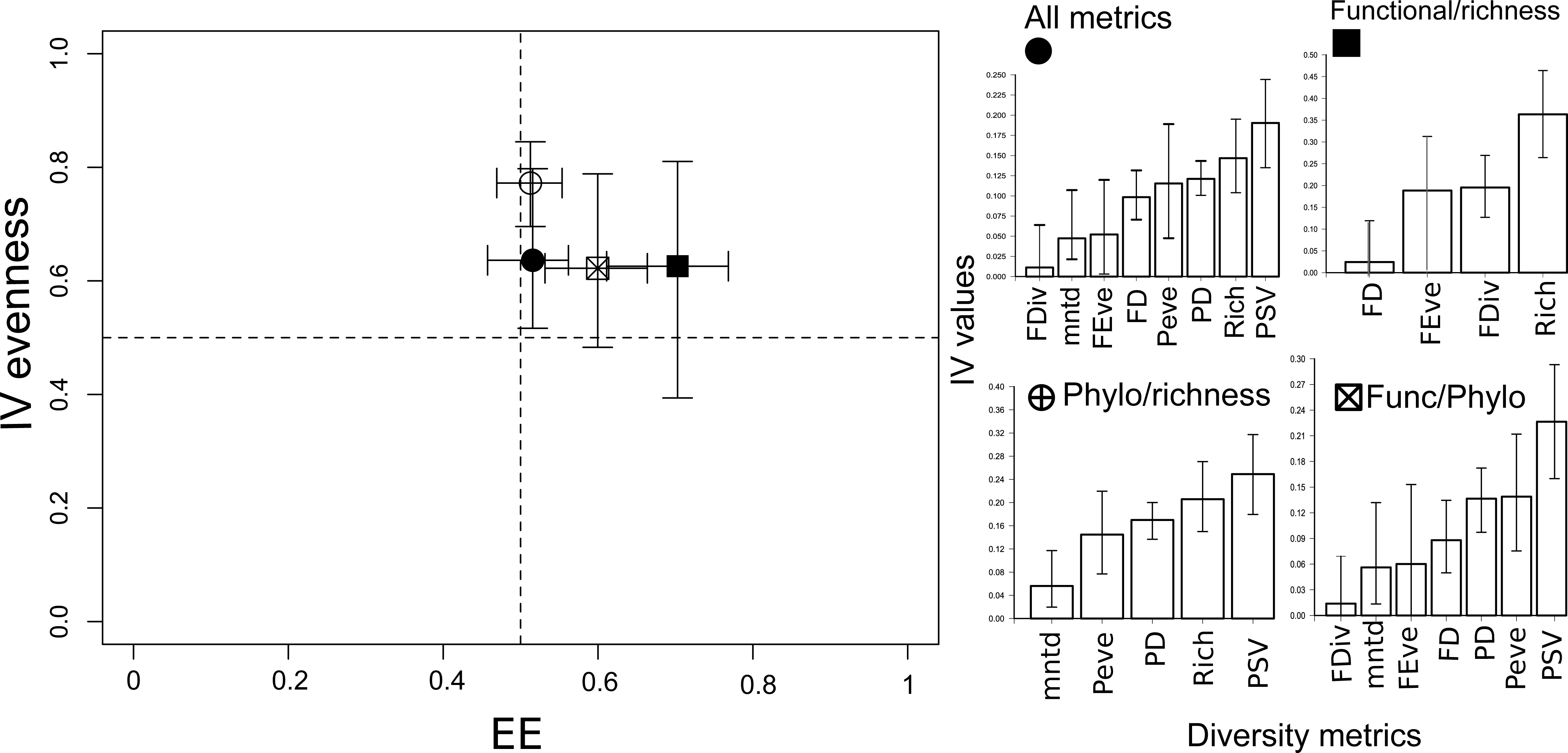
Values of EE and evenness of IV calculated for four different configurations of matrix **M**. Symbols represent mean values for each matrix configuration while lines represent confidence intervals. Bar graphics represent IV profiles calculated for matrix **M** with all metrics of diversity; functional metrics and richness; phylogenetic metrics and richness; and functional and phylogenetic metrics. Bars represent means while lines represent confidence intervals obtained via a bootstrap procedure applied to each matrix **M**.

## Discussion

Our results with simulated data evidence the need for a dimensionality framework that integrates both EE and IV in order to effectively characterize dimensionality by considering its two components —correlation and variation in biodiversity space. Operationalizing these two components through EE and IV reveals their complementarity (by means of EE) and, given some level of complementarity, the degree of redundancy in information captured by the metrics used to express these dimensions (through IV). Therefore, our proposed dimensionality framework represents a step beyond the current approach to operationalizing dimensionality, as proposed by Stevens and Tello (2014) by distinguishing the degree of redundancy in information that each diversity metric captures.

Our integrated dimensionality framework joins other propositions in helping to choose metrics for the biological characterization of communities. We are aware that the main guide for choosing diversity metrics must be the objectives of the work. However, regardless of the objective, it is desirable to use diversity metrics that encompass complementary components of biological diversity and account for a satisfactory amount of the information present in the biodiversity component being investigated (Ricotta 2005b). In this respect, Saito et al. (2015) showed that phylogenetic, functional and traditional taxonomic indices present complementary information and should be used to adequately characterize and monitor biodiversity of stream macroinvertebrate communities. Ouchi-Melo et al. (2018) performed an integrated assessment to identify areas of conservation interest in the Cerrado biome, and evidenced the importance of considering traditional together with functional and phylogenetic metrics. Although both of these works considered the complementarity component by accounting for correlation among metrics, they did not account for redundancy in the amount of variation that each metric captures in biodiversity space, thus facing the same problem presented by using the EE metric alone. The dimensionality framework presented here, therefore, represents the most general and complete framework to date for guiding researchers in their choice of metrics to be used for biological assessment by considering both complementarity among biological dimensions and the amount of information that metrics can capture.

It is worth pointing out that the dimensionality of diversity can be investigated at any spatial and temporal scale, and using any configuration of matrix **M**. Even for works that focus on only one component of biodiversity, the investigation of dimensionality can be important for knowing which aspects of biodiversity are worthy of being included in biological assessment. Tucker et al. (2017) identified three complementary components of the phylogenetic component: richness, divergence and regularity. Thus, research focused on phylogenetic diversity can address whether these three components are complementary dimensions in the analyzed communities and which metrics are the most important to measure in order to best represent variation in these dimensions. As we showed in our empirical example with small mammal communities, dimensionality will depend on the representation of biological diversity used in matrix **M**, which influences practical decisions regarding which metrics are the most important for characterizing biodiversity.

At least for the cricetid and marsupial communities analyzed here, characterizing diversity through functional and taxonomic components requires great care in the choice of diversity metrics to be used. This is because this situation has the highest complementarity regarding diversity dimensions (highest EE value), indicating the need to rely on different components of diversity to effectively describe biodiversity, and a moderate level of redundancy in metrics, indicating that some metrics account for disproportionately more information than others. In this example, richness accounted for more information than the other metrics, but consideration of other components that represent functional information is also important for effectively characterizing biological diversity. This functional component can be represented by FDiv or FEve, which are very redundant in information. On the other hand, if the characterization of small mammal communities was focused on phylogenetic and taxonomic components, the choice of metrics to be used would require less caution since complementarity among dimensions is lower and redundancy of information is greater, indicating that all the metrics capture similar amounts of information of biodiversity space.

When considering matrix **M** with all eight diversity metrics, applying the dimensionality framework to small mammal communities revealed that cricetids and marsupials possess intermediate to low levels of complementarity (mean EE of 0.51 ±0.025). Together with low complementarity, low levels of redundancy among the metrics was found when considering the three components of biodiversity together (mean IV evenness of 0.63 ±0.082). Consequently, we suggest that the choice of diversity metrics to effectively represent these communities must encompass the three components of diversity — choosing the PSV metric, which accounts for the highest IV, and two other complementary metrics to represent taxonomic (richness) and functional components (FD that has the highest IV among functional metrics, as shown in Figure 6).

The patterns of IV values for small mammal communities contrasted with the findings of Wilsey et al. (2005) and Lyashevska and Farnsworth (2012), who concluded that richness was the least important diversity metric for representing variation in community structure (grassland and marine benthic communities, respectively). Although we did not considered abundance-based metrics, as these studies did, we point out that patterns of complementarity and redundancy can differ depending on the taxonomic group being investigated and the metrics being used (as already emphasized by our empirical application of the IV framework with different configurations of matrix **M**). This finding highlights the need to understand contingencies in the correlation and variation components of the dimensionality of different communities.

We only used metrics that capture three sources of information from biodiversity (phylogenetic, functional and taxonomic), since they are the main assessed components of diversity and represent important metrics for capturing different dimensions of these components (Tucker et al. 2017). Despite the limited number of metrics presented in this work, the dimensionality framework used here is highly flexible in the sense that it can be applied to a matrix **M** that contains many more dimensions (Ricotta 2005). Therefore, we could represent diversity in a much more complete manner, with metrics that capture other quantifiable components such as genomic (*e*.*g*. Nei 1978), proteomic (*e*.*g*. Gotelli et al. 2013b) or any other dimension that can be quantified.

### Conclusion and future directions

This work represents an upgrade of the operationalization of the concept of dimensionality presented by previous works. We demonstrate that including the correlation component of dimensionality with the variation component, through the use of EE and IV, in the same framework more effectively characterizes the dimensionality of biodiversity.

Besides conceptual and operational advances, the dimensionality framework proposed here provides evidence regarding practical situations in which the choice of diversity metrics is more critical for effectively characterizing biodiversity. The use of this dimensionality framework can help identify these different situations and assist in choosing metrics.

Since the evidence presented in the literature regarding characterization of dimensionality is limited (Lyashevska and Farnsworth 2012, Stevens and Tello 2014, 2018, Stevens and Gavilanez 2015), and based only on specific groups of organisms, some questions still need to be addressed to provide a more complete understanding and generalization of the role that some factors play in the dimensionality of ecological communities. For instance, one might wonder if some dimensions of diversity are consistently more informative than others when describing diversity patterns among different taxa, or if distinct factors (historical, evolutionary and/or ecological) generate predictably higher or lower levels of dimensionality across communities.

## Acknowledgements

The authors thank W Ulrich, VD Pillar, V Debastiani and one anonymous review for invaluable suggestions in previous versions of the manuscript, and E Wild and E Bradley for proof-reading the English. GN received a PhD. Fellowship from Coordenação de Aperfeiçoamento de Pessoal de Nível Superior (CAPES). Research activities of LDSD was supported by Conselho Nacional de Pesquisa (CNPq) Productivity fellowships. GN and LDSD are members of the Instituto Nacional de Ciência e Tecnologia (INCT) in Ecology, Evolution and Biodiversity Conservation, supported by MCTIC/CNPq (proc. 465610/2014-5) and FAPEG.

## Notes

#### Summary of Updates

Introduction, Methods, Results and Discussion section was updated. We added Figure 5 to illustrate how dimensionality change depending on the configuration of matrix M.

https://gabrielnakamura.shinyapps.io/Supp_matShiny/

https://github.com/GabrielNakamura/dimensionality_function

https://github.com/GabrielNakamura/IV_function

