## Appendix 1 for "Revisiting the dimensionality of biological diversity"

Appendix S1 – Simulation procedure.

We created an interactive simulation module using the shiny package of RStudio, to allow the reader to explore the basic patterns and simulation procedures used in our work. The module contains a user interface (Figure S1 Section A) that presents the parameters used to simulate (A) phylogenetic trees, (B) species traits and (C) communities. These parameters are also used to calculate three diversity metrics (richness, phylogenetic diversity [Faith, 1999] and functional diversity [Petchey and Gaston, 2000]; sections E, F and G, respectively). Each modification that is made to any parameter of the user interface triggers the simulation procedure and generates a new phylogeny, new species traits, community composition and diversity metrics, following the same methods for simulating communities as described in the section ‘Simulation methodology*’* in the main text of this paper.

The default module begins by showing a pattern in which the community has a phylogeny containing 50 species (with a sliding bar called ‘Number of species in phylogeny’). The phylogenetic signal for species traits is high (set with a sliding bar called ‘Grafen parameter for phylogenetic signal’. Lower values of the Grafen parameter reduce phylogenetic signal, whereas higher values increase phylogenetic signal). The metacommunity was assembled by applying a phylogenetic habitat filter (*sensu* Duarte, 2002). At one extreme only the most phylogenetically related species (top 10%) can occur (set at the upper value of the sliding bar called ‘Lower and upper values of habitat filtering’), while at the other extreme of the phylogenetic gradient any species can occur (no phylogenetic restrictions, set at the lowest value of the sliding bar called ‘Lower and upper values of habitat filtering’). The phylogenetic filter create a gradient of phylogenetic and functional diversity (bar plots F and G). Richness can be set to be equal across all communities or a gradient can be applied by using the bottom widget (By selecting ‘Yes’ or ‘No’ in response to the question ‘Must richness be equal for all communities?’). Finally, the phylogenetic filter that is applied can be set to be either gradual (default) or not gradual (By selecting ‘Yes’ or ‘No’ in response to the question ‘Must the phylogenetic filter be gradual?’).

Figure S1: Layout of the interactive simulation module used to simulate metacommunities. The module comprises the basic parameters used for scenario simulation in the paper. A – buttons and sliding bars containing modifiable parameters that define the metacommunities; B – the phylogeny used to simulate traits and metacommunities and to calculate the PD metric; C – bar chart representing the trait values for each species in the phylogeny; D – representation of species incidence in the communities that compose the metacommunity, where the columns represent communities, lines represent species and the filled space represents the presence of a species in a community; E, F and G – bar plots representing the values for, respectively, richness (rich), phylogenetic diversity (PD) and functional diversity (FD) of each community.

Each time that a widget control is modified the simulation processes starts again by creating a new phylogeny, thus reflecting the same procedure used to perform simulations in the section ‘Simulation methodology’ of the main text. Using all possible combinations of widget controls will not produce all the scenarios showed in the main text; however, the widgets comprise the basic controls necessary to obtain the scenarios used. The application can be accessed at: https://gabrielnakamura.shinyapps.io/Supp_matShiny/.
