## Appendix 2 for "Revisiting the dimensionality of biological diversity"

Appendix S2 – Phylogenetic hypothesis and community data.

Figure S1 shows the phylogenetic hypothesis used to calculate the phylogenetic diversity indices used to assemble matrix **M**. The phylogenetic tree was assembled by using the mammal phylogenies of Bininda-Emonds et al. (2007) and Fabre et al. (2012). More details about the phylogenetic hypothesis used here can be found in the Methods section of the main text.

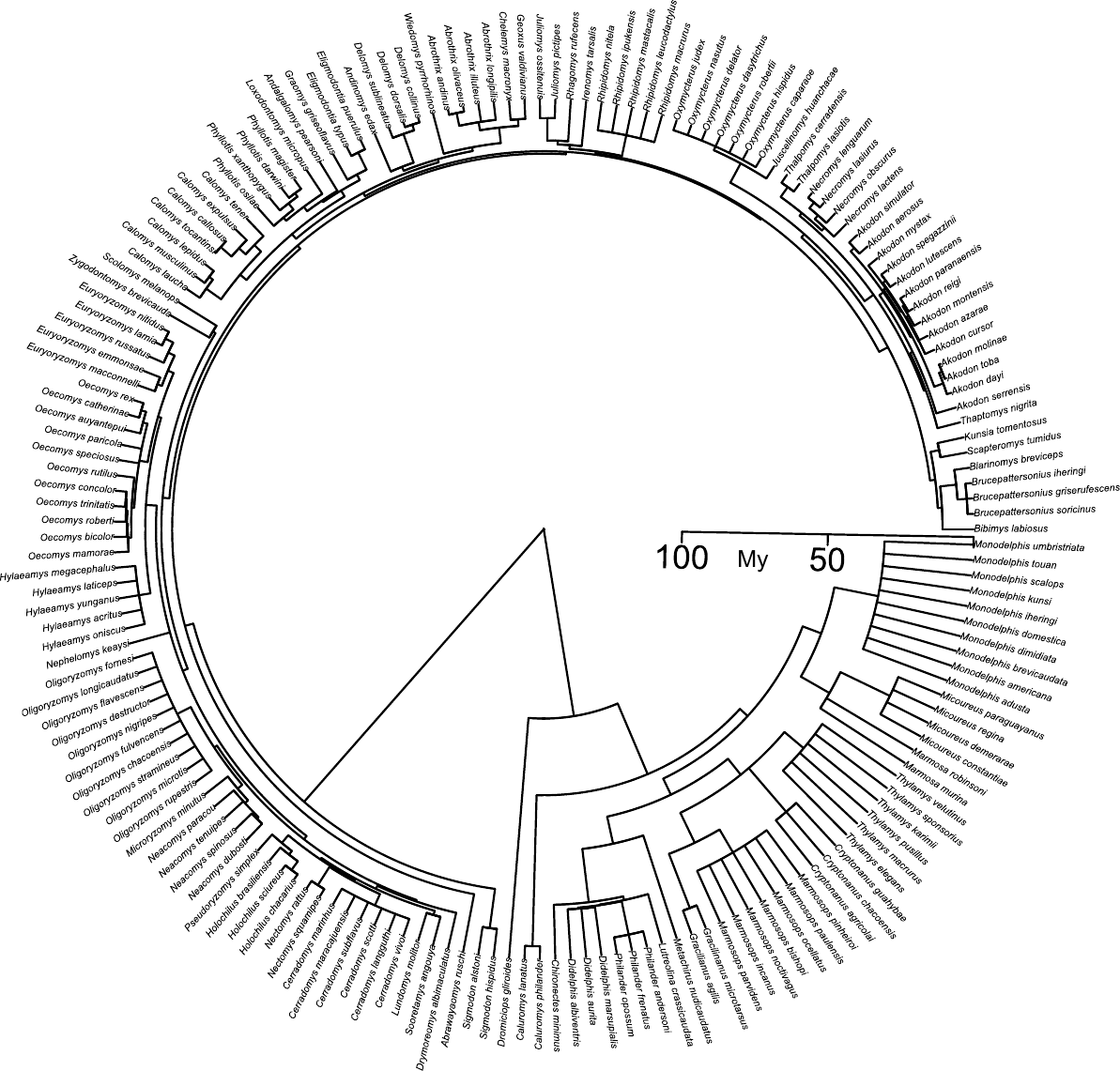

Figure S1: Phylogenetic hypothesis for small mammals (cricetids and marsupials) used to calculate phylogenetic metrics in this work.

Supplementary material – Appendix S2

Table S1: Reference, country, sampling effort (expressed as trap nights), latitude and longitude (decimal degrees) of the 103 small-mammal communities considered in this study.

| **Reference** | **Country** | **Trap-night** | **Latitude** | **Longitude** |
| --- | --- | --- | --- | --- |
| Kelt (2000) | Chile | 2894 | -40.50 | -73.00 |
| Thibault *et al.* (2011) | Argentina | 3416 | -39.92 | -71.42 |
| Saavedra & Simonetti (2004) | Chile | 8747 | -35.98 | -72.68 |
| Thibault *et al.* (2011) | Argentina | 1600 | -34.59 | -68.14 |
| Corbálan & Ojeda (2004) | Argentina | 4410 | -34.03 | -67.97 |
| Thibault et al. (2011) | Chile | 5528 | -33.38 | -70.52 |
| Munoz-Pedrero et al. (2010) | Chile | 1476 | -33.12 | -71.4 |
| Sponchiado *et al.* (2012) | Brazil | 4320 | -32.53 | -52.53 |
| Thibault et al. (2011) | Argentina | 36060 | -32.50 | -60.00 |
| Lima *et al.* (2010) | Brazil | 2240 | -29.67 | -53.72 |
| Marques et al. (2011) and Pedó et al. (2010) | Brazil | 5754 | -29.47 | -50.22 |
| Marques *et al.* (2011) | Brazil | 11596 | -29.42 | -50.40 |
| Dalmagro and Vieira (2005) | Brazil | 5178 | -29.17 | -50.08 |
| Graipel *et al.* (2006) | Brazil | 12132 | -27.72 | -48.53 |
| Cherem *et al.* (2011) | Brazil | 50097 | -27.63 | -48.83 |
| Ferro & Barquez (2009) | Argentina | 3440 | -27.20 | -65.93 |
| Melo *et al.* (2011) | Brazil | 6932 | -27.17 | -53.92 |
| Cirignoli *et al.*( 2011) | Argentina | 5310 | -27.1 | -54.97 |
| Ferro & Barquez (2009) | Argentina | 4181 | -26.67 | -65.63 |
| Quadros & Cáceres (2001) | Brazil | 1920 | -26.07 | -48.63 |
| Crespo (1982) | Argentina | >1000 | -25.52 | -54.13 |
| Quadros *et al.* (2000) | Brazil | 4380 | -25.48 | -53.12 |
| Bergallo *et al.* (1998) | Brazil | 5040 | -25.17 | -47.98 |
| Bergallo *et al.* (1998) | Brazil | 5040 | -24.53 | -47.25 |
| Vieira & Monteiro-Filho (2003) | Brazil | 3547 | -24.28 | -48.35 |
| Vieira & Monteiro-Filho (2003) | Brazil | 15227 | -24.23 | -48.06 |
| Pardini & Umetsu (2006) and Umestu & Pardini (2007) | Brazil | 9168 | -23.73 | -47.07 |
| Barros-Battesti *et al*. (2000) | Brazil | 2888 | -23.55 | -46.93 |
| Pinheiro & Geise (2008) | Brazil | 1680 | -23.37 | -44.83 |
| Bittencourt & Rocha (2003) | Brazil | 7474 | -23.18 | -44.20 |
| Jaksic *et al.* (1999) | Argentina | 2736 | -22.97 | -68.22 |
| Bonvicino *et al.* (2002) | Brazil | 1830 | -22.72 | -46.92 |
| Viveiros de Castro & Fernandez (2004) | Brazil | 51122 | -22.54 | -42.28 |
| Pires *et al.* 2002 | Brazil | 1618 | -22.52 | -42.28 |
| Vieira *et al.* 2009 | Brazil | 1200 | -22.5 | -42.86 |
| Geise *et al.* 2004 | Brazil | 1500 | -22.45 | -44.58 |
| Yahnke (2006) | Paraguyi | 23296 | -22.33 | -60.33 |
| Gentile *et al.* 2000 | Brazil | 12250 | -22.03 | -42.68 |
| Lyra-Jorge *et al.* (2001) | Brazil | 3672 | -21.62 | -47.62 |
| Talamoni & Dias (1999) | Brazil | 2400 | -21.55 | -47.85 |
| Cáceres *et al.* (2008) | Brazil | 1426 | -21.15 | -51.87 |
| Rocha *et al.* (2011b) | Brazil | 10080 | -20.88 | -44.83 |
| Paglia *et al.* (1995) | Brazil | 5760 | -20.75 | -42.85 |
| Cáceres *et al*. (2007) | Brazil | 430 | -20.7 | -56.85 |
| Moreira *et al*. (2009) | Brazil | 3168 | -20.67 | -42.43 |
| Thibault *et al*. (2011) | Brazil | 7264 | -20.52 | -41.00 |
| Cáceres *et al*. (2011) | Brazil | 11865 | -20.5 | -55.3 |
| Bonvicino *et al*. (2002) | Brazil | 3231 | -20.47 | -41.8 |
| Cáceres *et al.* (2010) and Hannibal & Cáceres (2010) | Brazil | 2950 | -20.45 | -55.53 |
| Pinto *et al.* (2009) | Brazil | 2160 | -20.37 | -40.47 |
| Passamani & Ribeiro (2009) and Passamani & Fernandez (2011) | Brazil | 23285 | -19.95 | -40.52 |
| Oliveira *et al*. (2007) | Brazil | 7115 | -19.93 | -43.88 |
| Andreazzi *et al.* (2011) | Brazil | 21560 | -19.88 | -56.38 |
| Grelle (2003) and Stallings *et al.* (1991) | Brazil | 68220 | -19.63 | -42.55 |
| Godoi *et al*. (2010) | Brazil | 4032 | -19.2 | -57.57 |
| Cáceres *et al*. (2011b) | Brazil | 16992 | -19.2 | -57.55 |
| Cáceres *et al*. (2011b) | Brazil | 4620 | -19.18 | -57.62 |
| Fonseca & Robinson (1990) | Brazil | 57120 | -19 | -42 |
| Rodrigues *et al*. (2002) | Brazil | 13562 | -18.25 | -52.88 |
| Cáceres *et al*. (2008) | Brazil | 1248 | -17.45 | -53.1 |
| Bonvicino *et al*. (1996) | Brazil | 1050 | -16.65 | -52.78 |
| Bonvicino *et al*. (1996) | Brazil | 2100 | -16.4 | -52.45 |
| Cáceres *et al*. (2008) | Brazil | 1350 | -16.37 | -51.95 |
| Aragona & Marinho-Filho (2009) | Brazil | 38635 | -16.23 | -56.37 |
| Thibault et al. (2011) | Brazil | 49810 | -16.07 | -47.92 |
| Santos & Henriques (2010) and Henriques *et al*. (2006) | Brazil | 2380 | -15.97 | -47.93 |
| Nitikman & Mares (1987) | Brazil | 12170 | -15.97 | -47.95 |
| Mares *et al*. (1986) | Brazil | 85101 | -15.93 | -47.88 |
| Santos & Henriques (2010) and Henriques *et al*. (2006) | Brazil | 3040 | -15.57 | -48.12 |
| Ribeiro e Marino-Filho (2005) | Brazil | 6600 | -15.53 | -47.6 |
| Santos-Filho et al. (2008) | Brazil | 17600 | -15.4 | -58.35 |
| Lacher Jr. & Alho (2001) | Brazil | 4821 | -15.33 | -55 |
| Pardini (2004) | Brazil | 7776 | -15.17 | -39.02 |
| Emmons (2009) | Bolivia | 13118 | -14.75 | -61.03 |
| Vargas & Simonetti (2004) | Bolivia | 4240 | -14.65 | -66.07 |
| Emmons (2009) | Bolivia | 6962 | -14.61 | -60.86 |
| Emmons (2009) | Bolivia | 7540 | -14.58 | -60.92 |
| Emmons (2009) | Bolivia | 4700 | -14.58 | -60.91 |
| Bonvicino et al. (2002) | Brazil | 2665 | -14.01 | -47.55 |
| Bonvicino et al. (2002) | Brazil | 1440 | -13.53 | -47.17 |
| Thibault et al*.* (2011) | Peru | 5600 | -13.14 | -69.61 |
| Thibault *et al*. (2011) | Peru | 1600 | -12.81 | -69.35 |
| Pereira & Geise *et al*. (2009) | Brazil | 10216 | -12.57 | -41.37 |
| Stevens & Husband (1998) | Brazil | 6144 | -11.18 | -37.42 |
| Bezerra *et al*. (2009) | Brazil | 5459 | -10.38 | -50 |
| Mena & Medellin (2010) | Peru | 5515 | -10.13 | -75.55 |
| Rocha *et al*. (2011) | Brazil | 32074 | -9.17 | -50.17 |
| Asfora & Pontes (2009) | Brazil | 1600 | -9 | -35.87 |
| Asfora & Pontes (2009) | Brazil | 1600 | -8.7 | -35.83 |
| Geise *et al*. (2011) | Brazil | 1391 | -8.53 | -37.25 |
| Asfora & Pontes (2009) | Brazil | 1600 | -8.25 | -35.08 |
| Thibault et al. (2011) | Brazil | 22800 | -7.77 | -51.96 |
| Thibault et al. (2011) | Peru | 5820 | -4.92 | -73.67 |
| Thibault et al. (2011) | Peru | 2888 | -4.50 | -73.42 |
| Hice & Schmidly (2002) | Peru | 2530 | -3.97 | -73.42 |
| Barnett & Cunha (1994) | Brazil | 3578 | -3.37 | -61.43 |
| Malcolm (1991) | Brazil | 25920 | -3.1 | -60.02 |
| Ribeiro Junior *et al*. (2011) | Brazil | 1206 | -1.95 | -51.60 |
| Thibault et al. (2011) | French Guiana | 4701 | 3.62 | -53.20 |
| Thibault *et al*. (2011) | Venezuela | 2225 | 7.40 | -62.82 |
| Thibault *et al*. (2011) | Venezuela | 38329 | 8.55 | -67.60 |
| Utrera *et al*. (2000) | Venezuela | 34455 | 9 | -69.75 |
| Thibault et al*.* (2011) | Venezuela | 1920 | 10.07 | -66.45 |
