## Appendix 3 for "Revisiting the dimensionality of biological diversity"

Appendix 3 – Additional information on simulation and empirical results for dimensionality framework presented in the paper.

We present here some additional results to support interpretations of the main results of simulation and empirical investigation (with small-mammal communities) of the dimensionality framework. Figure S3 is a boxplot containing F-values derived from two linear OLS models that described differences in IV among the three diversity metrics (PD, FD and richness) calculated for simulated communities with (A) high and (B) low correlation among diversity metrics.

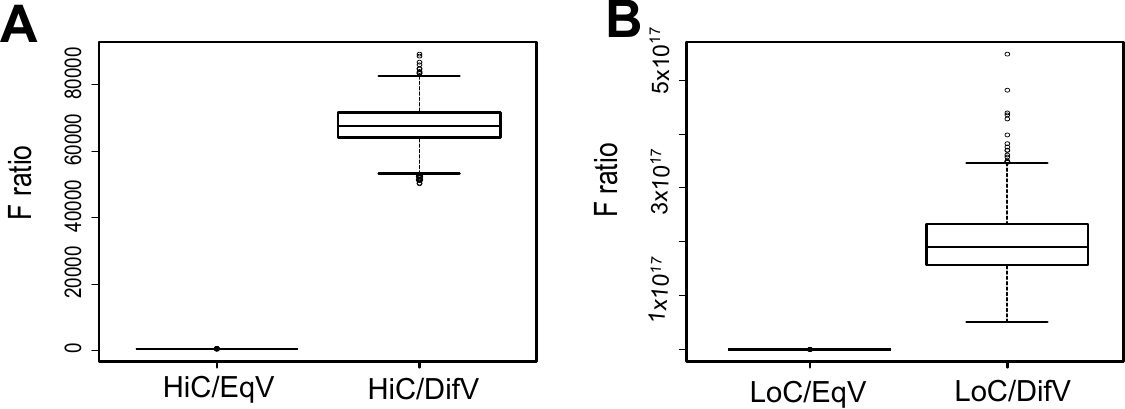

Figure S3: Boxplot with F-values for four simulated scenarios: HiC/EqV, HiC/DifV, LoC/EqV and LoC/DifV. Plot A shows F-values for a scenario with high correlation among metrics, while plot B shows F-values for scenarios with low correlation among metrics. The boxes represent the 25^th^, 50^th^ and 75^th^ quantiles.

We found that scenarios simulated with uneven variation among diversity metrics in biodiversity space had higher values for the F-ratio than scenarios with similar contribution, indicating great difference in IV among the three diversity metrics for both low and high correlation among metrics. It also becomes evident that differences among IV among the three diversity metrics were much higher for scenarios in which metrics were not correlated (boxplot B in Figure S3) than the scenarios of high correlation (boxplot A in Figure S3), indicating that the level of redundancy in metric importance can be much greater in scenarios with low complementarity (high correlation) than high complementarity (low correlation).

Figure S4 represents the null distribution and observed EE value calculated from matrix **M** containing all eight diversity metrics. The observed EE falls within the 95% confidence interval generated with a null model that changes species composition but keeps species richness of communities fixed.

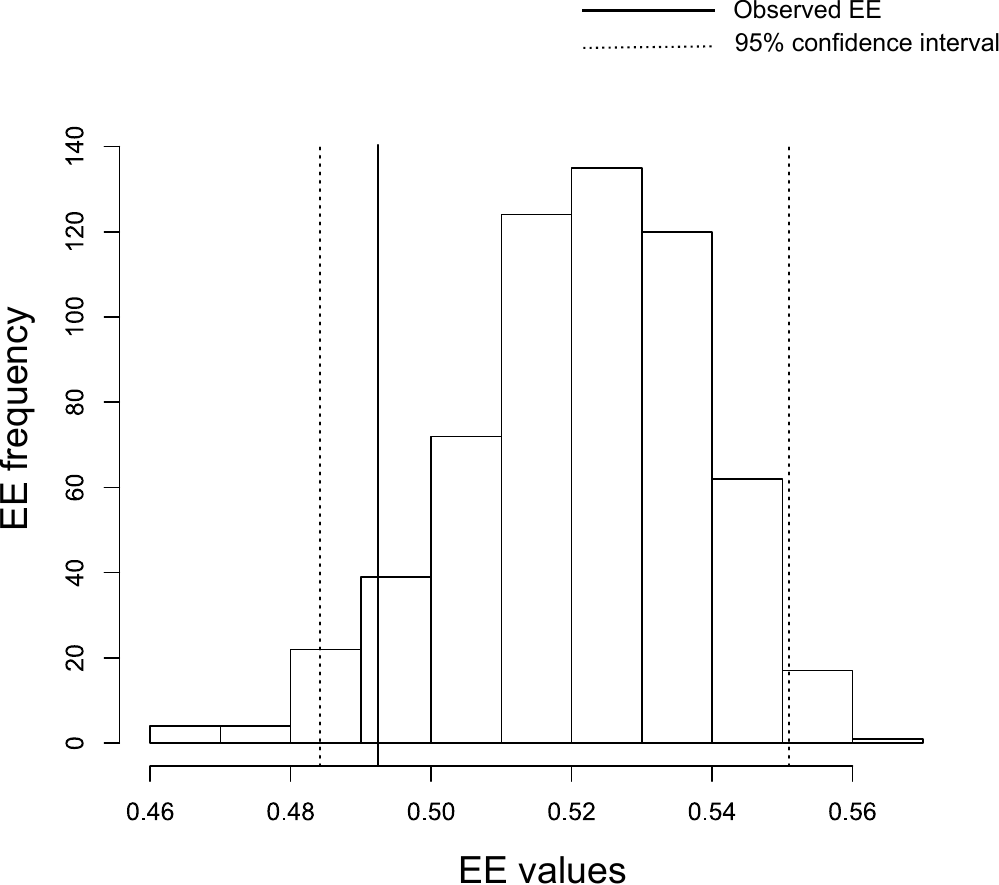

Figure S4: Null and observed values of EE for small mammal communities of cricetids and marsupials. The dotted lines represented the 95% confidence interval for a null model that changes community composition but keeps species richness fixed for each community. The continuous black line represent the observed EE calculated for matrix **M** with all eight diversity metrics.

We also performed Tukey tests using the two linear OLS models to compare IV values among the eight diversity metrics calculated for small mammal communities. Plot A shows pairwise comparisons among all diversity metrics, evidencing the greatest differences between PSV and functional indices. Plot B shows comparisons among the three components of diversity, which the eight indices represent, showing that the greatest difference in IV is between taxonomic and functional indices. This second Tukey test was performed by merging the metrics that represent a given component of diversity into the same category.

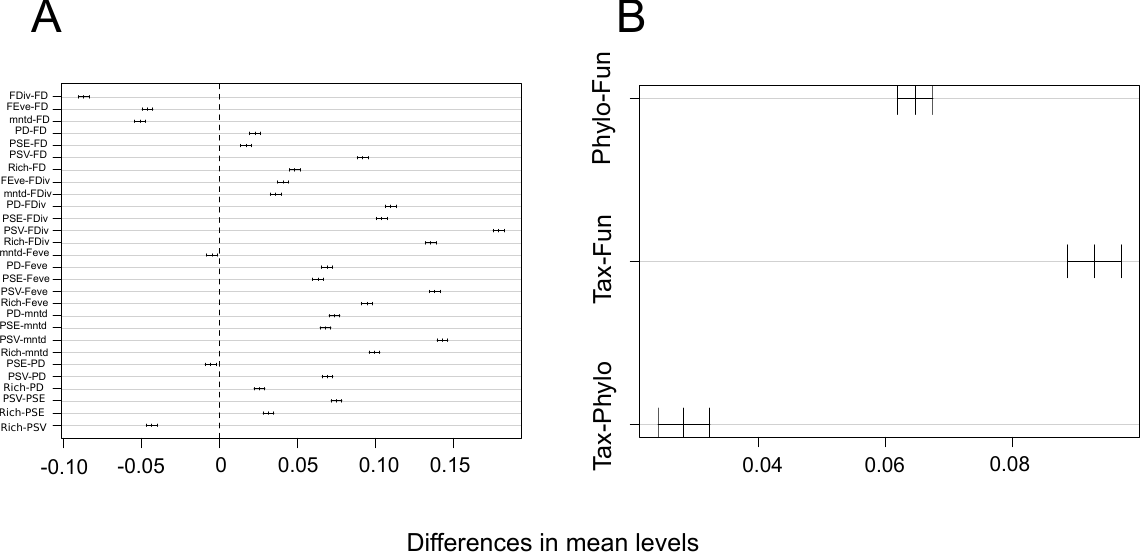

Figure S5: Plots showing mean differences for pairwise metric comparison (A) and pairwise comparison of functional, phylogenetic and taxonomic components of diversity (B). These two comparisons were obtained from a Tukey test of a linear OLS model.
